# Intravital imaging of osteocyte *α_v_β*_3_ integrin dynamics with locally injectable fluorescent nanoparticles

**DOI:** 10.1101/2023.02.23.529785

**Authors:** Melia Matthews, Emily Cook, Nada Naguib, Uli Wiesner, Karl Lewis

## Abstract

Osteocytes are the resident mechanosensory cells in bone. They are responsible for skeletal homeostasis and adaptation to mechanical cues. Integrin proteins play an prominent role in osteocyte mechanotransduction, however the details are not well stratified *in vivo*. Intravital imaging with multiphoton microscopy presents an opportunity to study molecular level mechanobiological events *in vivo*, and could be used to study integrin dynamics in osteocytes. However, fluorescent imaging limitations with respect to excessive optical scattering and low signal to noise ratio caused by mineralized bone matrix make such investigations non-trivial. Here we demonstrate that ultra-small and bright fluorescent core-shell silica nanoparticles (*<*7nm diameter), known as Cornell Prime Dots (C’Dots), are well-suited for the *in vivo* bone microenvironment and can improve intravital imaging capabilities. We report validation studies for C’Dots as a novel, locally injected *in vivo* osteocyte imaging tool for both non-specific cellular uptake and for targeting integrins. The pharmacokinetics of C’Dots reveal distinct sex differences in nanoparticle cycling and clearance in osteocytes, which represents a novel topic of study in bone biology. Integrin-targeted C’Dots were used to study osteocyte integrin dynamics. To the best of our knowledge, we report here the first evidence of osteocyte integrin endocytosis and recycling *in vivo*. Our results provide novel insights in osteocyte biology and will open up new lines of investigation that were previously unavailable *in vivo*.

## Introduction

Osteocytes are embedded in fully mineralized bone matrix and represent 90-95% of all cells found in bone tissue (1). Previously thought to be quiescent place-holders, it is now understood that they act as master conductors and regulators of skeletal health and homeostasis (2, 3). For their most important role, osteocytes are the resident mechanosensors in bone, incorporating mechanical signals along their processes and relaying biochemical signals through a highly interconnected dendritic network to matrix producing osteoblast cells and bone resorbing osteoclast cells (4–6).

*In vitro* work studying osteocyte mechanobiology over the last 25 years has established many details about osteocyte anatomy and mechanotransduction (7, 8). Osteocyte-like cell lines *in vitro* exhibit fluid flow induced activation leading to Ca^2+^ signaling (5) and bone anabolic cytokine expression (9). Osteocytes sense mechanical stimulation induced by fluid flow and generate biological response throughout their cellular network, stimulating mineral deposition and/or resorption (10). Additionally, it has been shown that osteocytes sense mechanical stimulation through a number of transmembrane proteins and channels along dendrites within the bone lacuno-canalicular space (LCS) (11). Integrin proteins are among those implicated as being key means of mechanical activation in osteocyte mechanotransduction *in vitro* (12–14). Indeed, McNamara and colleagues used electron microscopy to show physical evidence of integrin point attachments between osteocyte processes and their surrounding matrix (11), informing the updated model for mechanotransduction put forth by Qin et al. (15).

Integrins are heterodimeric proteins found within multiprotein complexes sensitive to mechanical stimulation on osteocytes called mechanosomes (12). Integrin mediated strain amplification is a key functional mechanism for osteocyte activation to mathematically predicted physiological fluid flow conditions (11, 14, 16). They are dynamic proteins involved in cell adhesion and mobility, and undergo endocytosis and recycling in numerous cell types (17–19). Two types of integrins exist in osteocytes, *α*_5_*β*_1_ and *α*_*v*_*β*_3_ (16, 20). The former is mainly expressed on the cell body while *α*_*v*_*β*_3_ integrins are heavily expressed on osteocyte processes (16). *α*_*v*_*β*_3_ integrins are known to undergo endocytosis as part of their role in cell migration and focal adhesion (21–23). Integrin dynamics involve clatherin-mediated cycling in osteoblast-like cells *in vitro* (24), but have not been previously observed in osteocytes.

Studying integrins *in vitro* has provided new information on the environment of osteocytes, but has limitations. For example, osteocyte mechanotransduction mechanisms are highly dependent on their native 3D microenvironment, which are difficult to recapitulate in 2D culture (20, 25, 26). One prominent manifestation of these limitations is that cells lying flat on a dish typically see polarized aggregation of integrins on the basal surface of the cell, removing them from typically apical side fluid activation (27). Recent advancements in imaging have made it possible to study osteocytes *in vivo*, providing an opportunity to expand our understanding of known osteocyte mechanotransduction mechanisms and dis-cover new details about their behavior in physiologically intact conditions.

Intravital imaging with multiphoton microscopy (MPM) is a powerful tool for studying osteocyte mechanobiology. Simultaneous absorption of multiple low energy photons of a laser beam focal point in MPM results in distinct imaging advantages, including reduced off-target excitation and a drastically higher signal to noise ratio at depth (28, 29). This has allowed for osteocytes and other bone cells to be visualized *in vivo* with genetically encoded fluorescent probes (30–32). One challenge with in vivo MPM imaging in bone is the dense optical scattering attenuation of traditional fluorophores, which has historically limited research questions to bulk cytosolic events such as Ca^2+^ signaling (31). Very few exogenous probes have been developed to target the subcellular dynamics of cells within bone, outside of Ca^2+^ signaling (33, 34). Research into a broader range of subcellular or molecular events *in vivo* has thus been restricted, and requires brighter more targetable fluorescent probes. Moreover, existing probes for intravital imaging are typically genetically encoded, requiring complex breeding strategies which are both expensive and time consuming. Locally injected, fast acting, and super bright fluorescent probes are needed to ask more complex and flexible questions about osteocytes *in vivo*.

Cornell Prime Dots (C’Dots) were developed for various bioimaging applications, both *in vitro* and *in vivo* (35–37). C’Dots are bright nanoparticles whose ultrasmall size (hydrodynamic diameter *<*7nm) permits access through the LCS to the area surrounding osteocyte cells (38). Their ultrasmall size leads to favorable biodistribution and pharmacokinetics profiles, proven in multiple human clinical trials with established biocompatibility (39, 40). Their silica core increases the brightness of encapsulated dyes 2.4 times via a combination of increases in radiative rates and decreases in nonradiative rates (35, 41–43). An additional beneficial aspect of C’Dots is that their brush-like poly(ethylene glycol) (PEG) shell (44), which minimizes protein corona formation and also enables facile covalent surface functionalization with various ligands (e.g., targeting or drug moieties), transforming them into targeted imaging or therapeutic probes (45–48).

Here we report the validation of C’Dots as an imaging tool for studying intracellular osteocyte dynamics *in vivo*. In addition, we use functionalized C’Dots with an RGD motif that targets integrins (49, 50). These RGD bearing C’Dots provide the first opportunity to interrogate integrin dynamics in osteocytes *in vivo*, with 3D microarchitecture, fluid environment, and endocrine interactions intact. Using this novel tool, we test the hypothesis that RGD functionalization increases the lifespan of C’Dots in osteocytes compared with non-functionalized C’Dots *in vivo*.

## Methods

### Animals

Male and female 16-20 week old C57Bl/6J mice (Jackson Laboratory, Bar Harbor, ME) were used in these experiments, representing skeletally mature young adult mice.

All procedures were approved by the Institutional Animal Care and Use Committees at Cornell University.

### Synthesis of PEGylated and cRGDyC Silica Core-Shell Nanoparticles

Ultrasmall fluorescent core-shell silica nanoparticles with encapsulated Cy5 dye (C’Dots) were synthesized by a modified Stöber process in water as previously described (35, 50). They were synthesized in two varieties: Untargeted control (PEG-C’Dots) and targeted (cRGDyC-C’Dots, labeled RGD-C’Dots in the rest of this manuscript). This synthesis approach involves the addition of tetramethyl orthosilicate (Sigma Aldrich) and silane-functionalized Cy5 organic dye (Lumiprobe) into a solution of water and ammonium hydroxide at a pH of 8 (35). The reaction was conducted at room temperature under vigorous stirring for 24 hours and was terminated by the addition of PEG-silane (EO6-9) (Gelest Inc). In the case of the cRGDyC C’Dots, cRGDy peptides (BioSynth Ltd.) containing the sequence cyclo-(Arg-Gly-Asp-Tyr) were conjugated to a heterobifunctional PEG-silane with a maleimido group through cysteine-maleimide linkage (Santa Cruz Biotechnology). This cRGDyC-PEG-silane was then added during the PEGylation step along with the unmodified PEG-silane (Gelest). Particles were then purified through gel permeation chromatography (GPC, BioRad Laboratories) prior to further optical characterization. Absorption and emission spectral profiles of the particles were obtained using a Varian Cary 5000 spectrophotometer (Varian, Palo Alto, CA) and a fluorescence spectrofluorometer (Photon Technology International). The hydrodynamic radius, particle brightness and sample concentrations were determined using a homebuilt fluorescence correlation spectroscopy (FCS) set-up using a 633 nm excitation solid state laser (35). As described in the Supporting Information, the PEG-C’Dots and cRGDyC-C’Dots exhibited similar properties including hydrodynamic radii (5.6 nm and 5.3 nm, respectively) and number of dyes per particle (2.0 and 2.1, respectively) as derived from a combination of FCS and UV-VIS data sets using methods detailed elsewhere (46). The number of cRGDyC peptides per C’Dot was estimated to be 20 ligands.

### C’Dot Injection and Incubation

Mice were anesthetized with 2-3% isoflurane mixed with medical grade air in an induction chamber for 3 minutes prior to injection. During injection, mice were kept under 2% isoflurane with a nose cone. A 15µL subcutaneous injection of C’Dots was then administered over the third metatarsal (MT3) in the hind paw (n=3-4/group/sex). Injection concentration was varied between 30µM and 10nM. In a separate experiment, incubation time after injection was varied from 5 minutes up to 24 hours with a fixed concentration of 10µM. Mice were allowed normal cage activity during their incubation time, with an exception for the 5 minute time point because it was too short to re-induce the mice after they awoke from the anesthesia.

### Metatarsal Isolation Surgery

Prior too surgery, anesthetized with 2-3% isoflurane mixed with medical grade air in an induction chamber for 3 minutes prior to injection. During MT3 isolation surgery, mice were kept under 2% isoflurane with a nose cone. A shallow vertical incision was made between the second and third metatarsal of the mouse hind paw and overlying tendons were removed. The MT3 was functionally isolated from the rest of the paw with a stainless steel pin beneath the mid-diaphysis of the bone, leaving primary vasculature at the proximal and distal palmar aspects intact. The bone was then stabilized in a 3 point bending configuration and submerged in room temperature DPBS prior to imaging, as reported previously (31). Mice were continuously anesthetized at 1.5-2% isoflurane during all subsequent imaging.

### C’Dot Imaging

C’Dot fluorescent signal inside the MT3 was visualized with multiphoton microscopy (MPM) (Bergamo II, Thorlabs). A 20x immersion objective (XLUMPLFLN, Olympus), 1090nm wavelength excitation, and a *>*647nm longpass filter acquisition was used. Images were captured at 1024×1024 pixel density and 0.588µm square pixels. To capture images of dendrites and subcellular C’Dot localization, 20 frames were averaged for a single image plane. For incubation time, concentration curve studies, and Calcein AM colocalization studies a 35µm z-stack was taken for each mouse per timepoint, starting 20µm beneath the bone surface, with 0.3µm steps between each frame and frame averaging of 7 frames per layer. For the clearance rate study, z-stacks„ were taken as described above every 15 minutes for almost 3 hours after the initial 1 hour incubation.

### C’Dot Image Analysis

Z-stacks were analyzed in ImageJ (NIH). A macro was created to interpolate the regions of interest across all layers of the z-stack as well as filter using despeckling and a Gaussian blur. Then 3D segmentation was run, creating objects for all cell volumes over a threshold intensity. The lower bound for threshold intensity was established as the mean plus 3x standard deviation of background fluorescence. Segmented objects were then filtered by surface area (*>*1000 pixels) to ensure only full cell volumes were counted, and then mean intensity for each object was quantified using the 3D manager tool.

### Intracellular C’Dot Validation

To evaluate intracellular localization of C’Dots, Calcein AM live cell stain (C-AM) was locally injected into the hind paw along with C’Dots (n=59 cells from 1 male mouse). The injection included 2.5µL of 100µM C-AM and 12.5µL of 10µM C’Dots. C’Dots were imaged using 1090nm wavelength excitation, and a *>*647nm longpass filter acquisition. C-AM was imaged at 920nm excitation and with a 490-560nm bandpass filter. Both stacks were loaded into ImageJ for analysis. Individual cells labeled with C’Dots or C-AM were identified manually and given their own ROI. Z-axis profiles of intensities were plotted and the range where signal increased from baseline into a spike was set as the bound of the signal. Percent spatial overlap of C’Dot and C-AM signal was quantified for analysis.

### Statistical Analyses

Differences between concentration groups and incubation time groups were assessed using a 2-way ANOVA followed by Tukey’s multiple comparisons test. Clearance study analysis was done using a Student’s t test and 2-way ANOVAs. Analyses were performed in GraphPad Prism version 9.0 for Mac OS X (GraphPad Software).

## Results

### Local Acute Delivery of C’Dots to Osteocytes *in vivo*

PEGylated Cornell prime dots (PEG-C’dots) covalently encapsulating Cy5 fluorescent dye were locally injected subcutaneously into the hind paw of C57Bl/6J mice to assess their potential as an *in vivo* fluorescent marker for osteocytes. Using MPM excitation at 1090nm (i.e., the near-infrared region), both collagen autofluorescence and C’Dot fluorescence were visualized in the MT3 diaphysis *in vivo* (**Figure 1**). C’Dot fluorescence was spatially diffuse through LCS in cortical bone, indicating C’Dots are able to travel through the pores in the LCS and interact with osteocytes. To assess uptake of the C’Dots into living osteocytes, Calcein-AM (a live cell stain) was co-injected with C’Dots and imaged. Robust co-localization of C’Dots with Calcein-AM was observed and almost 70% of C’Dot cell signal was fully contained within the C-AM signal (**Figure 2**).

**Fig. 1.**
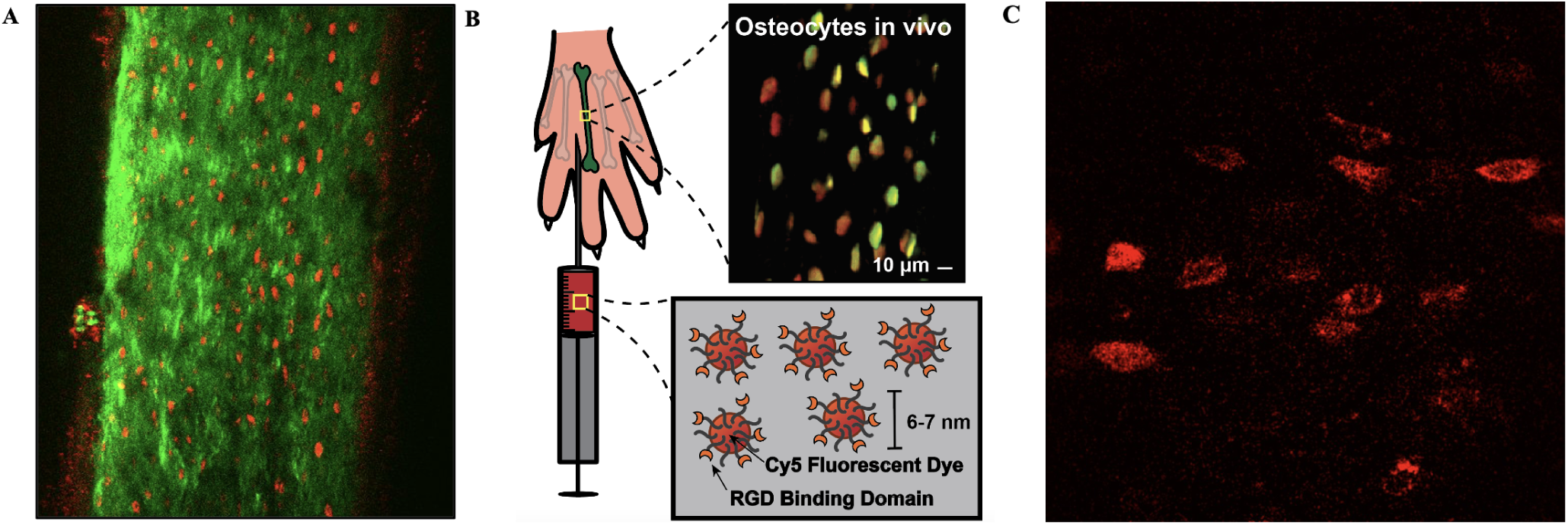
C’Dots allow successful intravital imaging of osteocytes in mice after a local injection above the MT3 bone. A) Representative MT3 MPM image showing autofluorescence of bone collagen (green) and Cy5 dye within C’Dots (10µM) (red). B) Localization of C’Dots within osteocytes visualized in a 35µM z-projection image. C) RDG-C’Dots visualized at 80x zoom reveal subcellular localization of C’Dots within osteocytes.

**Fig. 2.**
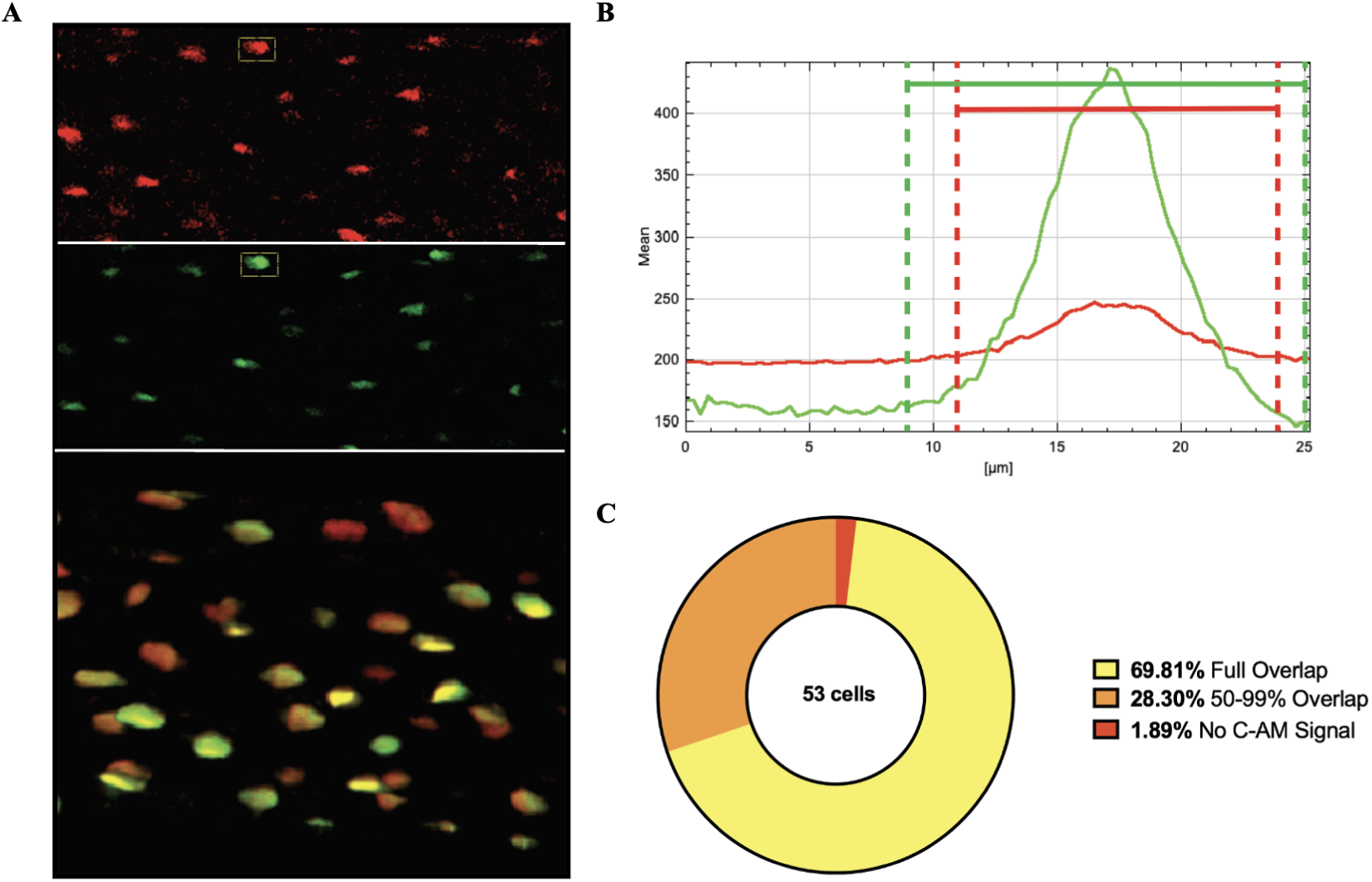
C’Dot co-localization with Calcein-AM *in vivo*. A) Representative image of C’Dot signal (red) and C-AM signal (green) with example cell regions of interest (dotted outline). 3D image overlay on the bottom. Channels were separated in ImageJ and individual cell ROIs were selected manually. B) Example z-stack intensity profile plot of single ROI in both channels. The range of cell signal was established as increase from baseline signal. C) Quantified C’Dot and C-AM overlap. 53 cell ROIs were quantified. Nearly 70% of C’Dot signal was fully contained within the range of C-AM signal. Only one ROI with C’Dot signal did not have C-AM signal as well. All ROIs with C’AM signal had over 50% overlap with the C’Dot signal. Average overlap was 94.8%, SD = ±8.9%.

### Non-Functionalized (Control) PEG-C’Dot Validation Experiments

Concentration dose-response curves and incubation time experiments were done to validate PEG-C’Dot use as a novel osteocyte probe in mouse metatarsal bone *in vivo*. The concentration of PEG-C’Dot injections were varied from 30µM down to 10nM, incorporating the range of PEG-C’Dot concentrations previously used in the literature for soft tissue imaging and adding higher concentrations to account for the dense, scattering material of bone (44). Strong but not fully saturated signal was seen down to 5µM, and became close to background fluorescence at 1µM and below (**Figure 3**). Reduction in signal with decreased C’Dot concentration was observed, as expected, and there were no significant differences between male and females at each concentration (**Figure 4A**). The best working concentration among those tested for PEG-C’Dots was found to be 10µM because there is strong signal with dynamic range available for both increases and decreases in signal. This ensures both detectable signal and dynamic range.

**Fig. 3.**
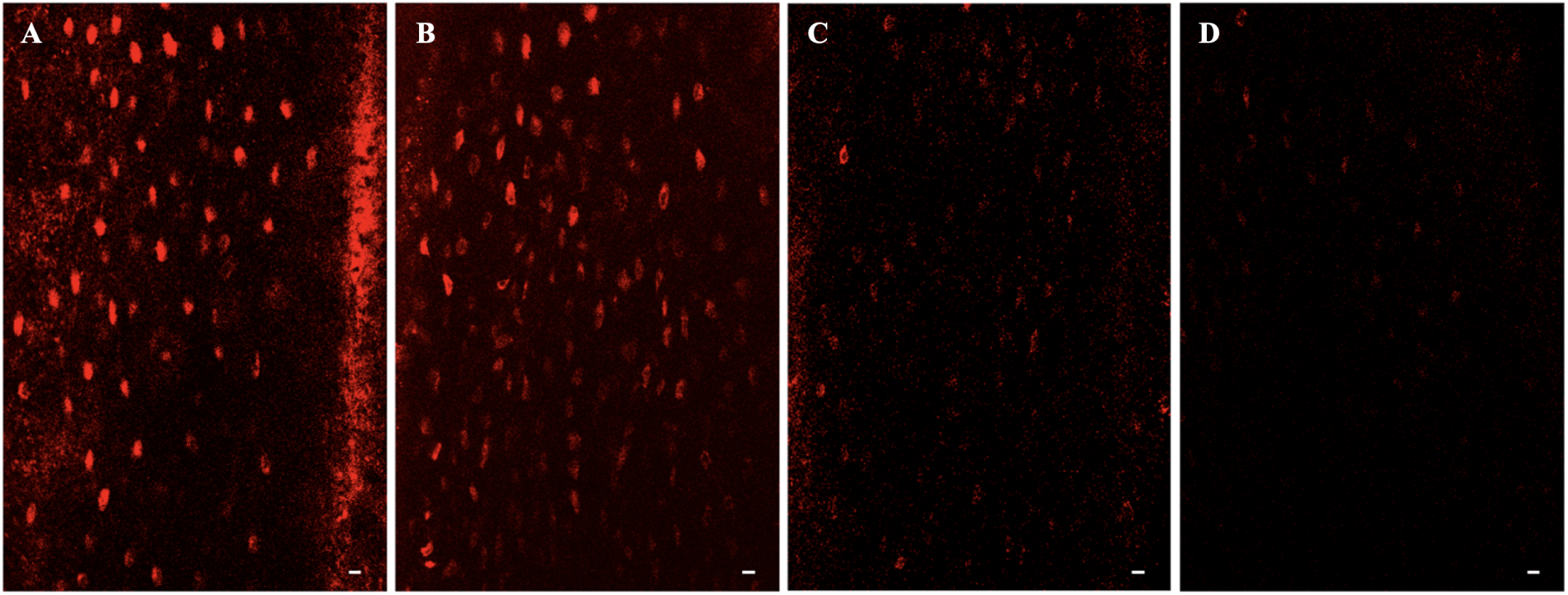
C’Dot fluorescence intensity decreases with subcutaneous injection concentration. Representative static images of MT3 diaphysis osteocytes after 1 hour of incubation using injection concentrations of A) 30µM, B) 10µM, C) 5µM, D) 1µM in male mice. C’Dots were excited at 1090nm and emission was acquired with a *>*647nm longpass filter. Scale bar = 10µM.

**Fig. 4.**
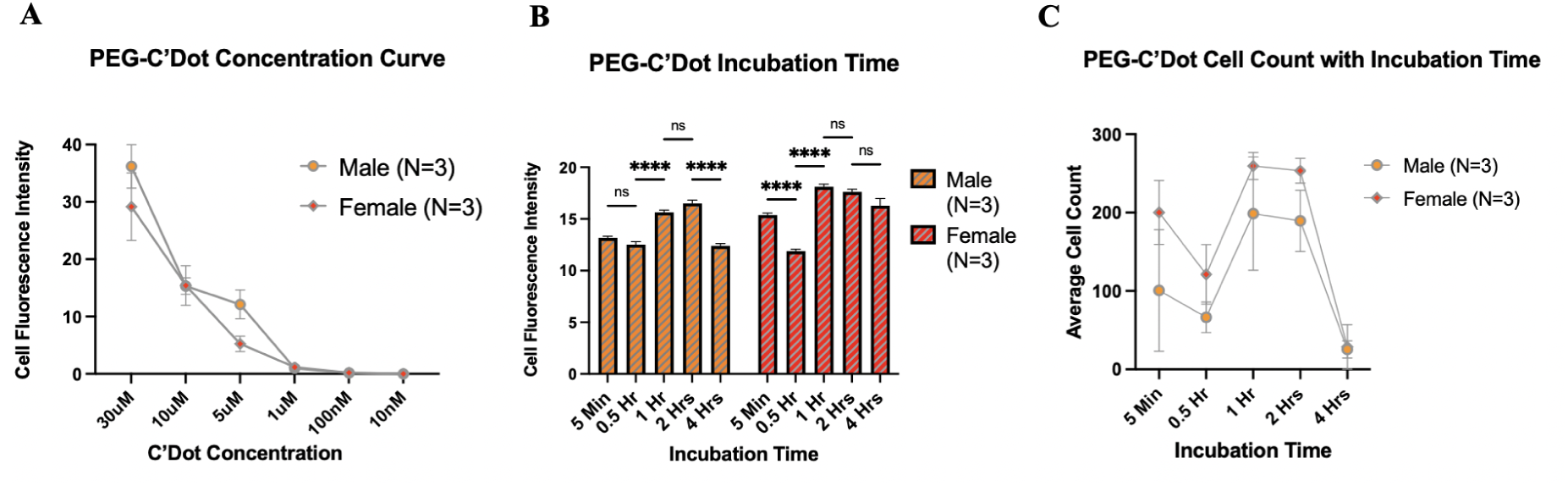
Experimental validation and optimization of non-functionalized PEG-C’Dots as a novel osteocyte imaging probe. A) Concentration dose-response intensity signal for PEG-C’Dots was lost at 1µM. Data points represent the average mean peak intensity for individual ‘cell’ ROIs. Significant differences were seen at distinct concentrations, but no significant differences were seen between groups with the same concentration. B) Incubation of PEG-C’Dots reveals distinct trends in 3D object intensity between male and female mice. Signal was retained out to 4 hours of incubation. Significant increases and decreases in intensity were observed, and the trends are distinct between the sexes. C) Cell count for male and female mice at each timepoint. Counts were averaged across all mice in the same experimental group and timepoint. \**p ≤* 0.05, error bars represent SEM, n=3 mice per group.

Holding the concentration steady at 10µM, incubation time was varied from 5 minutes out to 24 hours, with seven distinct timepoints (5min, 30min, 1, 2, 4, 12, and 24hrs). Fluorescence signal was quantifiable out to the 4hr incubation for both male and female groups (**Figure 4B**). Beyond those timepoints, the signal was too diffuse to be counted by 3D segmentation and filtering. Different fluctuation patterns in fluorescence intensity were observed between male and female mice, with peak increases occurring at different time points (**Fig 4B**). This result suggests unique dynamics between male and female osteocytes for the metabolism and clearance of PEG-C’Dots. Additionally, the 1hr timepoint had a consistent high intensity across the experimental groups, which will inform future experiments (**Figure 4C**).

### RGD Functionalized C’Dot Validation Experiments

Once proof of concept for PEG-C’Dot use as an *in vivo* osteocyte probe was established, and appropriate concentration and incubation times identified, targeted RGD-C’Dots were tested to see if the fluorescence intensity or dynamics would change. C’Dots functionalized with RGD motifs, which bind to *α*_5_*β*_1_ and *α*_*v*_*β*_3_ integrins, were characterized for *in vivo* osteocyte observation using the same methodology as the non-functionalized PEG-C’Dots (**Figure 1C**). It was hypothesized that RGD functionalization would allow C’Dot signal to be retained in the LCS/osteocyte for longer compared to non-targeted control PEG-C’Dots due to targeting and binding of RGD-C’Dots to integrins, which are not cleared from the LCS under any known mechanisms. Concentration dose-response curve and incubation time were again assessed using the same groupings as the PEG-C’Dot studies (**Figure 5A,B**). Signal was retained at slightly lower concentrations for RGD-C’Dots compared to PEG-C’Dots, above background fluorescence out to 100nM instead of 1µM. The preferred experimental concentration was consistent with PEG-C’Dots at 10µM. Signal was once again lost beyond 4hrs of incubation time, confirming fast RDG-C’Dot clearance. Incubation of 1 hour retained high intensity signal across groups (**Figure 5C**). The temporal intensity trends seen in PEG-C’Dot incubation time experiments switched between males and females when using the RGD-C’Dots. Specifically, the strong drop in signal at the 30min incubation timepoint and subsequent rebound at the 1hr timepoint seen in control PEG-C’Dot females was now being seen in the RGD-C’Dot males. This suggests potential differences in integrin and C’Dot endocytosis and recycling depending on sex.

**Fig. 5.**
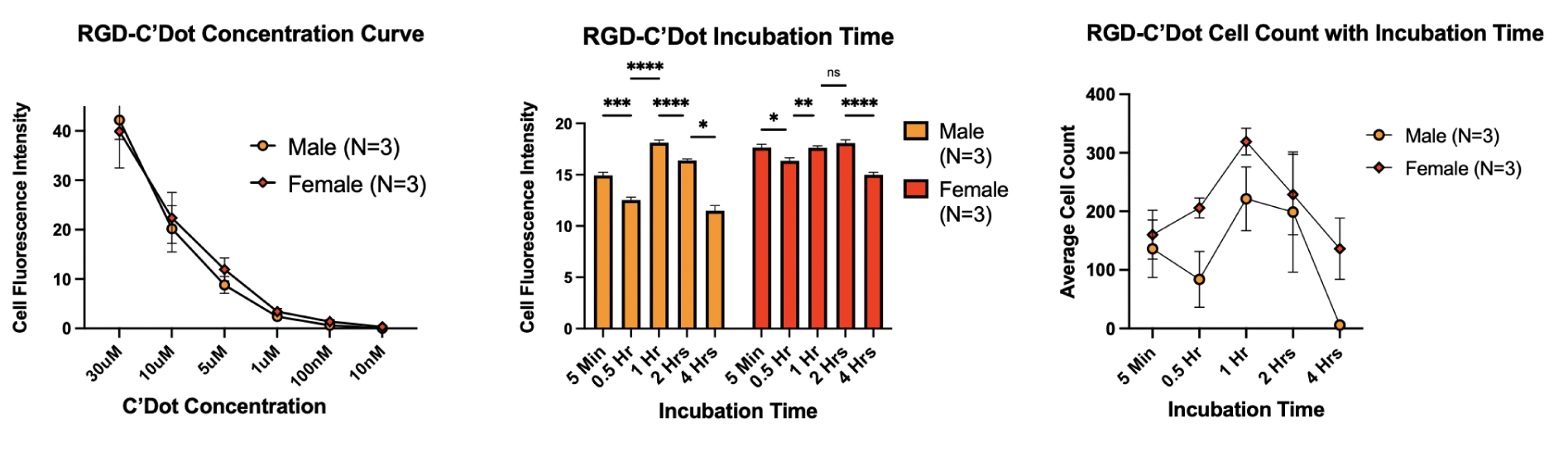
Validation and optimization of functionalized RGD-C’Dots use as a osteocyte integrin targeted probe. A) Concentration dose-response intensity signal for RGD-C’Dots was lost at 100nM, a lower concentration than the control C’Dots. Optimal sub-saturation signal was again selected at 10µM. Data points represent the average mean peak intensity for individual ‘cell’ ROIs. Significant differences were seen at distinct concentrations, but as with control C’Dots, no significant differences were seen between groups with the same concentration. B) RGD-C’Dot intensity based on incubation time was distinct from the dynamics observed in control C’Dots, and had a reversal in biphasic trend between the sexes. Signal was again lost after 4 hours. C) Cell count for male and female mice at each timepoint. Values were averaged across mice. \**p ≤* 0.05, error bars represent SEM, n=3 mice per group.

### Clearance Rate of Functionalized RGD-C’Dots and Non-Functionalized PEG-C’Dots from LCS Over Time

After establishing concentration and incubation time in control PEG-C’Dots and targeted RGD-C’Dots, the dynamics of both groups of C’Dots were studied within the LCS over time in individual mice. This was done to better understand the pharmacokinetics of the C’Dots and to set a baseline measurement for serial PEG-C’Dot and RGD-C’Dot imaging. The number of cells that were quantified decreased markedly over time and changed in distinct ways between the sexes. Male mice, using either targeted RGD-C’Dots or control PEG-C’Dots, had total loss of signal after 1hr of imaging (**Figure 6A**). Female mice, on the other hand, retained at least some signal to the end of the experiment, displaying an unmistakable difference in trend line as compared to males (**Figure 6A**). Male mice clearing C’Dots from the LCS more quickly than their female counterparts suggests sexual dimorphism in fluid flow and/or osteocyte C’Dot metabolism. Male mice also exhibited differences between PEG and RGD-C’Dots in measured intensity over time, with PEG-C’Dot cell volumes having lower average volumetric intensity compared to RGD-C’Dots while females did not (**Figure 6B,C**). Of note, C’Dot intensity in males could no longer be accurately quantified after 90 minutes compared to 150 minute in the females (**Figure 6B,C**). Time lapse images of C’Dot clearance from the LCS over time was visualized using a side-by-side video format comparing males to females depicts these differences dynamically (data not shown). For both control PEG-C’Dots and targeted RGD-C’Dots, it is clear that male mice lost signal much earlier compared to time-matched females.

**Fig. 6.**
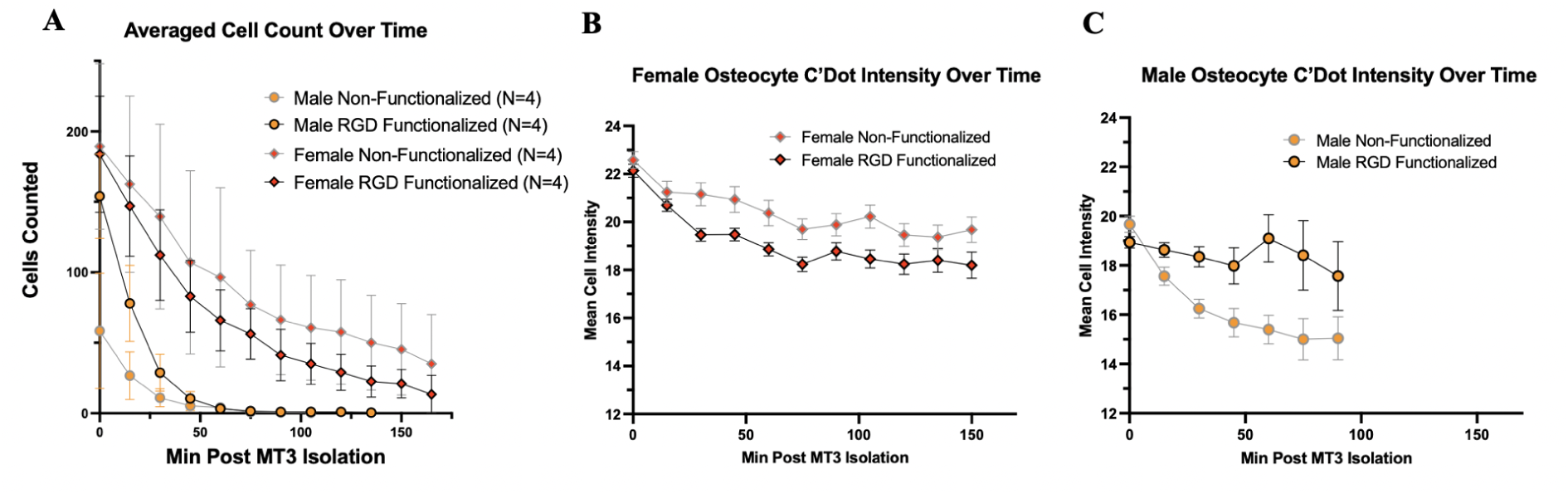
Analysis of imaged non-functionalized PEG-C’Dot and functionalized RGD-C’Dot z-stacks over time reveals unique clearance between male and female mice. A) Number of 3D ‘cells’ were counted over time, out to almost 3 hours post MT3 isolation. Z-stacks were taken 15 minutes apart using an automated script in Thor software. 3D objects or ‘cells’ were quantified from each z-stack using an ImageJ macro. Male signal is negligible after one hour, while female signal remains out to the end of the experiment. Targeted RGD-C’Dots and Control PEG-C’Dot groups are not significantly different within each sex. N= 3 or 4 mice per group. B,C) Mean intensity for each counted cell was averaged within experimental group at each timepoint. Between 1 and 700 cells were averaged for each data point. Female RGD and Control were significantly different at one timepoint. ** *p ≤* 0.01. Error bars = SEM.

## Discussion

Osteocytes were successfully observed *in vivo* in wild-type C57BL/6 mice in a timeframe on the order of minutes to hours using MPM and locally injected exogenous fluorescent probes, C’Dots. C’Dots provide a novel opportunity to interrogate osteocytes *in vivo*, with implication for anatomical and functional study applications, including rapid observation of live cell intracellular dynamics. The current paucity of bone cell-specific exogenous fluorescent probes along with the time and monetary expense of genetically modified fluorescent mouse lines make C’Dots a very useful improvement in the realm of intravital bone cell imaging (28).

Co-localization of C’Dot fluroescent signal with Calcein AM (**Figure 2**) suggests they are being absorbed into osteocyte cytosol, likely via endocytosis. Our data show that the C’Dot signal is taken up by live osteocytes in almost every instance, albeit to varying degrees. The finding of nanoparticle endocytosis is supported by the fluorescent dynamics of both control PEG and RGD-C’Dots, which suggest that endocytosis results in clearance or degradation of the C’Dots over time. C’Dots have been shown to undergo endocytosis in other cell and tissue types, but their half-life in osteocytes and the LCS is remarkably short by comparison. Benezra et al. observed that C’Dot signal was retained in tissue for multiple days (51), while signal from osteocytes within the LCS strongly dissipated only hours after injection. That the C’Dots clear from osteocytes much faster than other cell and tissue types is intriguing; it implies rapid nanoparticle trafficking and, by extension, very active intracelluar osteocyte shuttling of small objects (e.g., proteins, foreign bodies, cellular membrane components, etc). This type of trafficking may be due to endocytosis of C’Dots followed by intracellular degradation. Alternatively, endocytosis may be followed by recycling to the cell-surface and clearance via interstitial fluid. Although C’Dots have a strong silica coating, osteocytes have been shown to have low pH capabilities used in localized bone mineral resorption (52, 53). They may be able to degrade the endocytosed C’Dots with their highly active ATPase proton pumps. However, it is currently unknown if the C’Dots are being actively degraded or ejected into circulation, and this mechanism should be further stratified.

Endocytosis and cycling of C’Dots was also suggested by the validation experiment for incubation time. Unexpectedly, male and female mice were observed to have different timelines of change in intracellular fluorescence dynamics following C’Dot injection, with females exhibiting more variability in intensity during the first three timepoints measured. This trend was then seen in males when using the integrin targeting RGD-C’Dots. This difference between C’Dot dynamics in males and females suggests that functionalization of the C’Dots and binding to integrins has somehow altered their metabolism and clearance within the LCS in a sex dependent manner. The differences between control PEG-C’Dots and targeted RGD-C’Dots support potential of cycling of RDG-C’Dot bound-integrins within osteocytes *in vivo*. Integrin cycling has been shown in other bone cells (24), but has not previously been reported in osteocytes. Based on their prominent role in mechanobiology, understanding the dynamics of integrins *in vivo* will be a valuable tool in studying cellular accommodation, desensitization of osteocytes to mechanical loading, and osteocyte metabolism and intracellular trafficking.

Distinct trends in fluorescent signal were observed between male and female mice as a function of clearance rates. Females retained signal for much longer compared to males. Female PEG-C’Dots also exhibited the strongest intensity over time and male control PEG-C’Dots showed the lowest. Male control mice also lost signal 30 minutes prior to RDG-C’Dots mice. This may align with our original hypothesis that functionalization can allow for longer signal retention, however it is curious that it is only observed in male mice. That C’Dot clearance from the LCS appears to be sexually dimorphic was an unexpected finding, but not without precedent. Sex differences in osteocyte protein expression (54) and sex hormones are known to have sizeable impacts on bone health and homeostasis (55, 56). Those sex dependent mechanisms may also have a role in the membrane dynamics of bone cells. Sex differences in endocytosis of membrane proteins such as integrins represents an interesting field of research for understanding sexual dimorphism in osteocyte function and could be a potential target for minimizing bone loss in aging, especially for post-menopausal women. For example, bisphosphonates such as incadronate are potent anti-resorption drugs, and they are also known to inhibit endocytosis (57, 58).

Overall, our results prove the use of ultrasmall and bright particle probes like C’Dots as potent *in vivo* osteocyte probes, likely able to reveal details of endocytosis and membrane dynamics in osteocytes, which to the best of our knowledge has not been previously observed *in vivo*. Additionally, sexual dimorphism in osteocyte metabolism and C’Dot clearance was observed, potentially providing a novel area of research to target for bone diseases. Future studies will confirm endocytosis of PEG-C’Dots and their targeted analogue, i.e., RGD-C’Dots, localize their movement over short timeframes to track intravital integrin metabolism in osteocytes, and interrogate sex-specific differences in osteocyte dynamics.

## ACKNOWLEDGEMENTS

We thank Karly Hooper for maintaining mouse colonies for use in these experiments, along with the entire CARE staff in Weill Hall. This work was supported by the Department of Biomedical Engineering at Cornell University.

## Supplementary Information

**Fig. 7. Figure S1:**
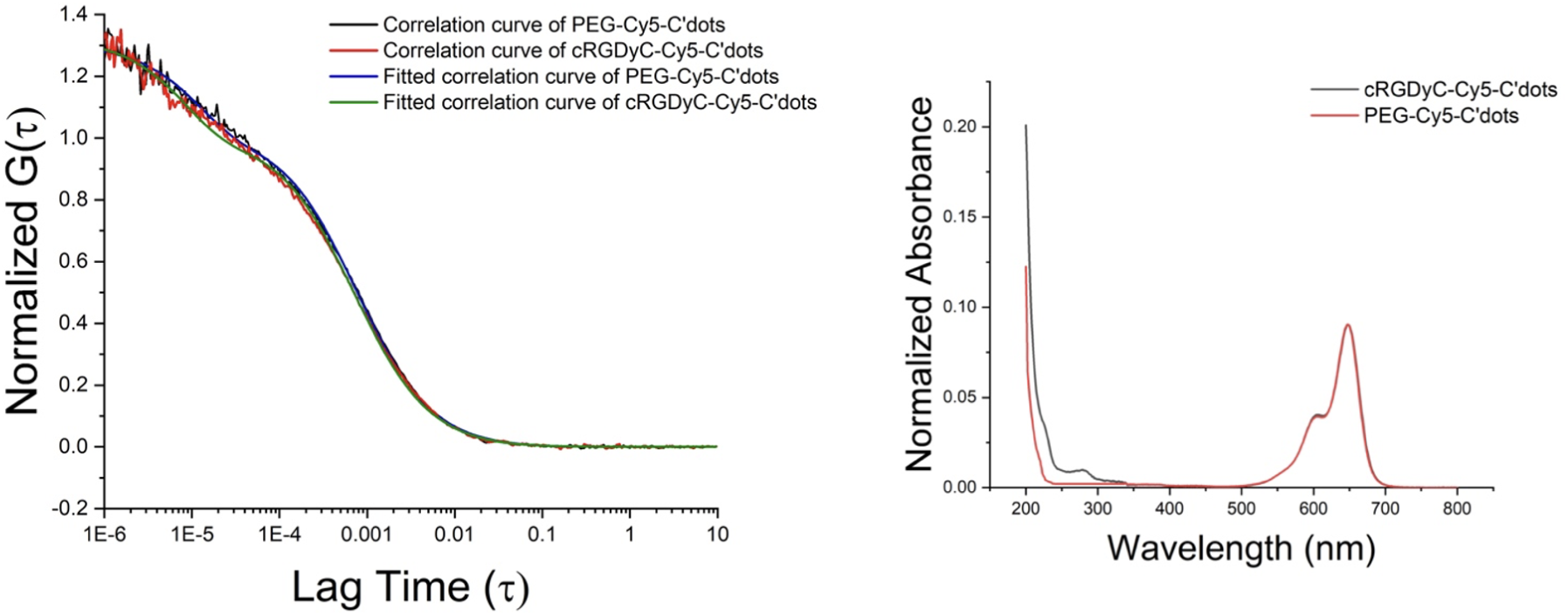
Characterization of PEG-Cy5-C’dots and c(RGDyC)-Cy5-C’dots. (Left) FCS correlation curves and fits of purified PEG-Cy5-C’dots and c(RGDyC)-Cy5-C’dots, suggesting hydrodynamic diameters of 5.6 nm and 5.3 nm, respectively. (Right) Comparison of UV-VIS spectra of PEG-Cy5-C’dots without (red) and after functionalization with c(RGDyC) (black). The absorbance peak at 275 nm corresponds to that of c(RGDyC) and suggests around 20 RGD ligands per particle.

