## Supplemental Figures for "Intravital imaging of osteocyte *α_v_β*_3_ integrin dynamics with locally injectable fluorescent nanoparticles"

### Supplementary Information

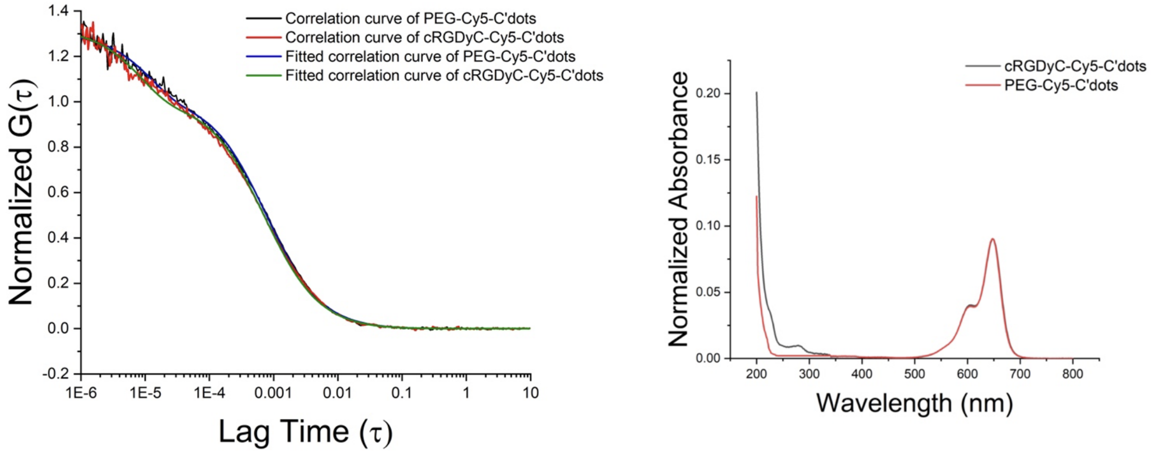


**Fig. 7.** Figure S1: Characterization of PEG-Cy5-C'dots and c(RGDyC)-Cy5-C'dots. (Left) FCS correlation curves and fits of purified PEG-Cy5-C'dots and c(RGDyC)-Cy5-C'dots, suggesting hydrodynamic diameters of 5.6 nm and 5.3 nm, respectively. (Right) Comparison of UV-VIS spectra of PEG-Cy5-C'dots without (red) and after functionalization with c(RGDyC) (black). The absorbance peak at 275 nm corresponds to that of c(RGDyC) and suggests around 20 RGD ligands per particle.
